# Proficiency Testing for bacterial whole genome sequencing in assuring the quality of microbiology diagnostics in clinical and public health laboratories

**DOI:** 10.1101/2020.09.18.304519

**Authors:** Katherine A. Lau, Anders Gonçalves da Silva, Torsten Theis, Joanna Gray, Susan A Ballard, William D. Rawlinson

## Abstract

The adoption of whole genome sequencing (WGS) data over the past decade for pathogen surveillance, and decision-making for infectious diseases has rapidly transformed the landscape of clinical microbiology and public health. However, for successful transition to routine use of these techniques, it is crucial to ensure the WGS data generated meet defined quality standards for pathogen identification, typing, antimicrobial resistance detection and surveillance. Further, the ongoing development of these standards will ensure that the bioinformatic processes are capable of accurately identifying and characterising organisms of interest, and thereby facilitate the integration of WGS into routine clinical and public health laboratory setting. A pilot proficiency testing (PT) program for WGS of infectious agents was developed to facilitate widely applicable standardisation and benchmarking standards for WGS across a range of laboratories. The PT participating laboratories were required to generate WGS data from two bacterial isolates, and submit the raw data for independent bioinformatics analysis, as well as analyse the data with their own processes and answer relevant questions about the data. Overall, laboratories used a diverse range of bioinformatics tools and could generate and analyse high-quality data, either meeting or exceeding the minimum requirements. This pilot has provided valuable insight into the current state of genomics in clinical microbiology and public health laboratories across Australia. It will provide a baseline guide for the standardisation of WGS and enable the development of a PT program that allows an ongoing performance benchmark for accreditation of WGS-based test processes.

## INTRODUCTION

The portability, reproducibility and potential anonymity of whole genome sequencing (WGS) data, along with its ability to provide high-resolution comparisons both within and across jurisdictions, represent an important advance and have become the underlying forces driving the transition to genomics in public health. The higher throughput, and exponential increase in the quality and flexibility, as well as the rapidly decreased cost and turnaround time mean that the next-generation sequencing (NGS) technologies are rapidly becoming a viable approach in complementing or even replacing existing molecular technologies and conventional assays currently run simultaneously in a diagnostic microbiology laboratory [1], [2].

However, to fully integrate NGS into clinical and public health laboratory settings, we need to first address issues including establishment of quality assessment (QA) and quality control (QC) measures, and the development of practice guidelines for accreditation purposes in ensuring the quality of NGS-based tests. It is crucial to ensure that the WGS data generated meet minimum quality standards and that the bioinformatic processes are capable of accurately identifying and characterising the organisms of interest. Acting as an external quality assessment (EQA) tool, proficiency testing (PT) is not only useful for evaluation and verification of sequencing quality and reliability in bioinformatic analyses, it also independently assesses the test performance of clinical laboratories to ensure the quality, harmonization, comparability, and reproducibility of diagnostic results.

Instead of the traditional analyte-specific PT, the assessment of interlaboratory performance recommended for the clinical NGS application is based on the method used, due to the extremely large variety of possible target sequences. This assessment can be achieved based on the test results performed on blinded samples provided to participating laboratories. In the absence of a formal PT program, sample exchange with a laboratory performing similar tests can be utilized as an alternative assessment activity [3]. Since 2015, the Global Microbial Identifier (GMI) has been the forerunner in offering annual EQA and PT for WGS applications of infectious agents https://www.globalmicrobialidentifier.org/workgroups/about-the-gmi-proficiency-tests [4]. Other similar PT has been established for U.S. Food and Drug Association (FDA) field laboratories to focus on food pathogens [5], [6], while a more recent global PT was offered to assess bioinformatics analyses of simulated *in silico* clinical WGS data of virus in improving the identification of emerging diseases [7]. More recently, the CDC has released guidelines to assist laboratories adopt NGS workflows [8], and a global consortium, PH4AGE (https://pha4ge.github.io/) has been established to help translate many of the standards developed by the GA4GH (https://www.ga4gh.org/) to the microbial world.

To facilitate the standardisation and benchmarking of WGS across a range of laboratories, the Pilot PT program for WGS of infectious agents was developed and offered in November 2018. This pilot PT was designed to be applicable to laboratories that use a variety of sequencing platforms and test applications. In this two-part PT, participating laboratories were first required to generate WGS data from two *Salmonella enterica* subsp. *enterica* isolates, and along with their no-template control (NTC), submit the raw data for independent bioinformatics analysis. The second part focussed on the bioinformatics pipeline of participating laboratories, and required laboratories to analyse the WGS data with their processes and provide further information about the data and their bioinformatics pipeline. The WGS data submitted by participating laboratories in this study were assessed against a set of minimum data quality standards and bioinformatic processes, which were specifically established for this pilot PT. A workflow of standards and a fully reproducible bioinformatics analysis pipeline were made available so that participating laboratories could replicate the exact analysis carried out with their WGS data.

Overall, the pilot PT program has provided significant understanding into the current state of transition of genomics into Australian public health. Participation in a WGS PT program will support clinical and public health laboratories in the implementation of WGS as a diagnostic and epidemiological tool.

## MATERIALS AND METHOD

### Organisation and participating laboratories

The pilot PT program was initiated by the Royal College of Pathologists of Australasia Quality Assurance Programs (RCPAQAP), in collaboration with the Communicable Diseases Genomics Network (CDGN), an Expert Reference Panel under the Public Health Laboratory Network (PHLN), Australia. This was based on the relevant information collected from an initial online questionnaire (Supplemental file) sent to participating laboratories of RCPAQAP Microbiology, RCPAQAP Serology and RCPAQAP Molecular Infectious Diseases in February 2018. The questionnaire was provided in four sections with the aims to understand and capture i) the basic details of end-users; ii) the capabilities of the laboratory in participating in a WGS pilot PT program; iii) the most frequently targeted infectious agents; and iv) the current QA practice at the laboratory. Following this, the PT program was announced through email invitation to a total of 10 Australian clinical and public health laboratories experienced in analysing WGS data sets for infectious agents.

### Survey specimens and survey instructions

Participating laboratories were supplied with two samples consisting of live bacteria isolates from the genus *Salmonella*, grown on chocolate agar slopes. Characteristics of these samples are recorded in Table 1.

**TABLE 1.** Characteristics of the samples sent to the participating laboratories

| **RCPAQAP**<br>**SEQID** | **Genus, species,**<br>**subspecies** | **Serotype** | **Sequence type** | **Notes** |
| --- | --- | --- | --- | --- |
| BSQAP001 | *Salmonella*<br>*enterica* subsp.<br>*enterica* | Enteritidis | ST3304 | Clade A |
| BSQAP005 | *Salmonella*<br>*enterica* subsp.<br>*enterica* | Typhimurium | ST34 | ser 4,[5],12:i:- |

Participating laboratories were instructed to perform WGS on both isolates, and an NTC according to their established protocols, and report results within a six-week timeframe (from 8 November to 19 December 2018). The laboratories were to analyse the sequence data using their choice of bioinformatics software (commercial or in-house pipeline) and i) to identify the genus, species (and sub-species, if applicable) of the organisms; ii) to report on the *in silico* multilocus sequence type (MLST) of the organisms, if a scheme exists; iii) to report on antimicrobial resistance gene(s) detected; and iv) to provide the *in silico* serotype of the organisms, if appropriate for the organisms.

### Submitted PT results

Results generated in this pilot PT program were captured using two reporting methods. Firstly, the laboratories were to upload and submit the raw FASTQ data set for the two cultures and an NTC to a dedicated, secure File Transfer Protocol (FTP) server site within a six-week timeframe (from 8 November to 19 December 2018). Participating laboratories were provided with a unique laboratory identifier (LABID) to ensure that the overall results were anonymized.

Secondly, participating laboratories were instructed to report all results obtained using their own bioinformatics analysis pipeline(s) via an online survey link (https://www.surveymonkey.com/) (see Table S1 in the supplemental material). The survey contained five sections, including the identification of the specimens, a report on the protocol used to generate the WGS data, the criteria used to identify the species, the criteria used to evaluate the quality of the sequence data, and the bioinformatics software and tool used for analysis. Laboratories were also invited to provide any feedback for the survey. The responses were collected as single or multiple options with free text for remarks and comments.

### Analysis of PT results

The definition of the minimum data quality standards and the development of a bioinformatics pipeline used to analyse the data were initiated following the discussion within the CDGN Bioinformatics Working Group. The PT results for both samples were assessed based on six quality metrics; i) the average quality score (Q-score); ii) the inferred genome coverage; iii) the inferred organism identification; iv) the genome assembly size; v) the inferred MLST scheme; and vi) the inferred MLST sequence type (ST), while the NTC was assessed only based on the total sequenced nucleotides; the number of sequenced nucleotides for NTC must be less than 1,000,000 base pairs (bp) to pass this assessment. The definition of the criteria for quality metrics used to assess the performance of participating laboratories for BSQAP001 and BSQAP005 is available in Table S2 in the supplemental material.

The FASTQ data were analysed by the CDGN Bioinformatics Working group. The bioinformatics pipeline engine used for these analyses was implemented using Snakemake [9]. All the bioinformatic tools were installed in Singularity containers [10]. All tools were installed using the package manager Conda (https://docs.conda.io/en/latest/), and both version and build were specified (see Table S3 in the supplemental material). In short, read statistics were generated with *seqtk* [11]; reads were trimmed with *trimmomatic* [12] using a database of Illumina adapter sequences; species identification was performed using *kraken2* [13] with the MiniKraken Database v2 (includes bacteria, Archaea, virus, and human genomes) released on 2018-11-01; genome size and coverage estimation from *k*-mers was performed using *mash* [14]; read assembly was performed using *shovill* [15] using the *SPAdes* assembler [16]; assessment of the draft assembly was done with *quast* [17]; MLST was inferred using *mlst* [18] using schemes available on PubMLST [19]; presence of antimicrobial resistance (AMR) genes was inferred using *abricate* [20] using the NCBI AMR reference database [21]; and serotype was inferred using *SISTR* [22], a tool for inferring the serotype of *Salmonella enterica*.

All containers are stored in CloudStor and available for download (see Table S4 in the supplemental material). Finally, the whole process can be reproduced using *vagrant* (https://www.vagrantup.com/) and *virtualbox* (https://www.virtualbox.org/) following the instructions available on the GitHub repository (https://www.github.com/cdgn-anz/wgsptp-pipeline). The repository contains all the code for the pipelines and recipes for building the Singularity containers. Two Snakemake pipelines were written and named: *data* (to process the FASTQ data associated with each sample) and *ntc* (to process the samples labelled as “negative control”); and both used the same elements.

### Pilot PT report

The report issued to the participating laboratories had two parts. Part one contained the summary of the identification of both samples, as well as the details of the protocol used to generate the WGS data, while part two contained the individual report on the QC performance of each laboratories. The comparative data for BSQAP001 and BSQAP005 based on a total of six data aspects (average Q score, percent guanine-cytosine (GC), total nucleotides, total reads, assembly N50 and assembly size) derived from the analysis performed on the raw data set submitted by all participating laboratories were compared based on the Z-scores of each aspect. The Z-score was determined by subtracting the mean from the data point, and this value was then divided by the standard deviation to normalise the data. The observation of the variation in Z-scores of different data aspects were plotted on the same graph. This provided laboratories with a quick comparison across all aspects of the data across laboratories.

## RESULTS

### Identification (ID) of pilot PT samples

A total of eight laboratories identified the species in both samples using a *k*-mer ID approach, while two laboratories adopted a 16S rRNA gene ID approach for species identification (Table 2). The details of the list of top three species hits identified in the sequence data for BSQAP001 and BSQAP005, including the percentage (%) of reads that matched these hits or the evidence and confidence of each hit, are available in Table 2. The criteria used by laboratories with a *k*-mer ID approach to identify the species and the criteria used to evaluate the quality of the sequence are available in Table S5 and Table S6 in the supplemental material, respectively.

**TABLE 2.** Top three species hits identified for; A) BSQAP001, B) BSQAP005, using kmer and 16S identification (ID) approaches

| ID approach | Participating laboratory | Species<sup>a</sup> |  |  |  |  |  |
| --- | --- | --- | --- | --- | --- | --- | --- |
|  |  | Species 1 |  | Species 2 |  | Species 3 |  |
| Kmer | 1 | *Salmonella enterica* | 97.04% | *Salmonella enterica* subsp. *enterica* | 95.87% | *Salmonella enterica* subsp. *enterica* serovar Dublin | 1.44% |
|  | 2 | *Salmonella enterica* | 82.78% | *Escherichia coli* | 0.43% | *Salmonella bongori*; | 0.01% |
|  | 3 | *Salmonella enterica* | 93.86% | *Escherichia coli* | 1.12% | *Enterobacter aerogenes* | 0.02% |
|  | 4 | *Salmonella enterica* | 93.15% | *Escherichia coli* | 1.09% | *Klebsiella pneumoniae* | 0.02% |
|  | 5 | *Salmonella enterica* |  | unclassified |  | *Burkholderia dolosa* |  |
|  | 6 | *Salmonella enterica* | 93.29% | *Salmonella enterica* subsp. *enterica* | 88.80% | *Salmonella enterica* subsp. *enterica* serovar Dublin | 0.59% |
|  | 7 | *enterica* | 96.99% |  |  |  |  |
|  | 10 | *Salmonella enterica* | 84.56% | *Escherichia coli* | 0.59% | *Enterobacter aerogenes* | 0.02% |
| 16S | 8 | *Salmonella enterica* |  | *Salmonella enterica* |  | *Salmonella enterica* |  |
|  | 9 | CP030214.1 *Salmonella enterica* strain SA20025921 | 99% | CP016357.1 *Salmonella enterica* subsp. *enterica* serovar Newport str. WA_14882 | 99% | CP010284.1 *Salmonella enterica* subsp. *enterica* serovar Newport str. CVM N1543 | 99% |

B. BSQAP005
| ID approach | Participating laboratory | Species<sup>a</sup> |  |  |  |  |  |
| --- | --- | --- | --- | --- | --- | --- | --- |
|  |  | Species 1 |  | Species 2 |  | Species 3 |  |
| Kmer | 1 | *Salmonella enterica* | 93.73% | *Salmonella enterica* subsp. *enterica* | 92.71% | *Salmonella enterica* subsp. *enterica* serovar Typhimurium | 1.58% |
|  | 2 | *Salmonella enterica* | 80.93% | *Escherichia coli* | 0.15% | *Erwinia pyrifoliae* | 0.13% |
|  | 3 | *Salmonella enterica* | 90.69% | *Escherichia coli* | 0.68% | *Edwardsiella ictaluri* | 0.58% |
|  | 4 | *Salmonella enterica* | 90.58% | *Citrobacter freundii* | 0.28% | *Escherichia coli* | 0.16% |
|  | 5 | *Salmonella enterica* |  | unclassified |  | *Burkholderia dolosa* |  |
|  | 6 | *Salmonella enterica* | 88.89% | *Salmonella enterica* subsp. *enterica* | 84.82% | *Salmonella enterica* subsp. *enterica* serovar Typhimurium | 0.72% |
|  | 7 | *enterica* | 93.09% |  |  |  |  |
|  | 10 | *Salmonella enterica* | 93.12% | *Escherichia coli* | 0.43% | *Edwardsiella ictaluri* |  |
| 16S | 8 | *Salmonella enterica* |  | *Salmonella enterica* |  | *Salmonella enterica* |  |
|  | 9 | CP019649.1 *Salmonella enterica* subsp. *enterica* serovar Typhimurium strain TW-Stm6 | 99% | CP029568.1 *Salmonella enterica* strain DA34837 | 99% | LN999997.1 *Salmonella enterica* subsp. *enterica* serovar Typhimurium isolate SO4698-09 | 99% |

Results are presented relative to the 10 participating laboratories. All laboratories correctly identified BSQAP001 and BSQAP005, respectively, as *Salmonella enterica* subspecies *enterica*. In regards to serotyping BSQAP001, six laboratories reported the sample to be Enteritidis, two reported the serogroup (D1), and two reported the antigenic formula (1,9,12:g.m:-). In sample BSQAP005 we observed a little more variation. A total of three laboratories reported the sample as Typhimurium, four laboratories reported the antigenic formula (I 4,[5],12:i:-), and the remaining three laboratories reported either serogroup B), or the (potential) monophasic variant of Typhimurium. Details in Fig. 1. All but one laboratory reported the correct *in silico* MLST for BSQAP001 (ST3304) and BSQAP005 (ST34) (Fig. 2). Regarding the detection of AMR genes, we observed substantial variation in the reported genes harboured by both samples (Table 3). Despite no AMR gene matches identified for BSQAP001 based on the analysis performed by the Bioinformatics Working Group, CDGN on all raw FASTQ data set submitted by participating laboratories, there were six laboratories who identified aac(6’)-Iaa, three laboratories that reported mdf(A), and only four laboratories reported the absence of any AMR genes. A total of five AMR genes (tet(B), sul2, blaTEM-1, aph(6)-Id, aph(3”)-Ib), either complete or partial matches were identified for BSQAP005, based on the analysis performed on all raw FASTQ data set submitted, by eight laboratories. Laboratories also reported four other AMR gene matches, aac(6’)-laa (n=5), mdf(A) (n=), strB_1 (n=1) and strA_4 (n=1),.

**FIG 1:**
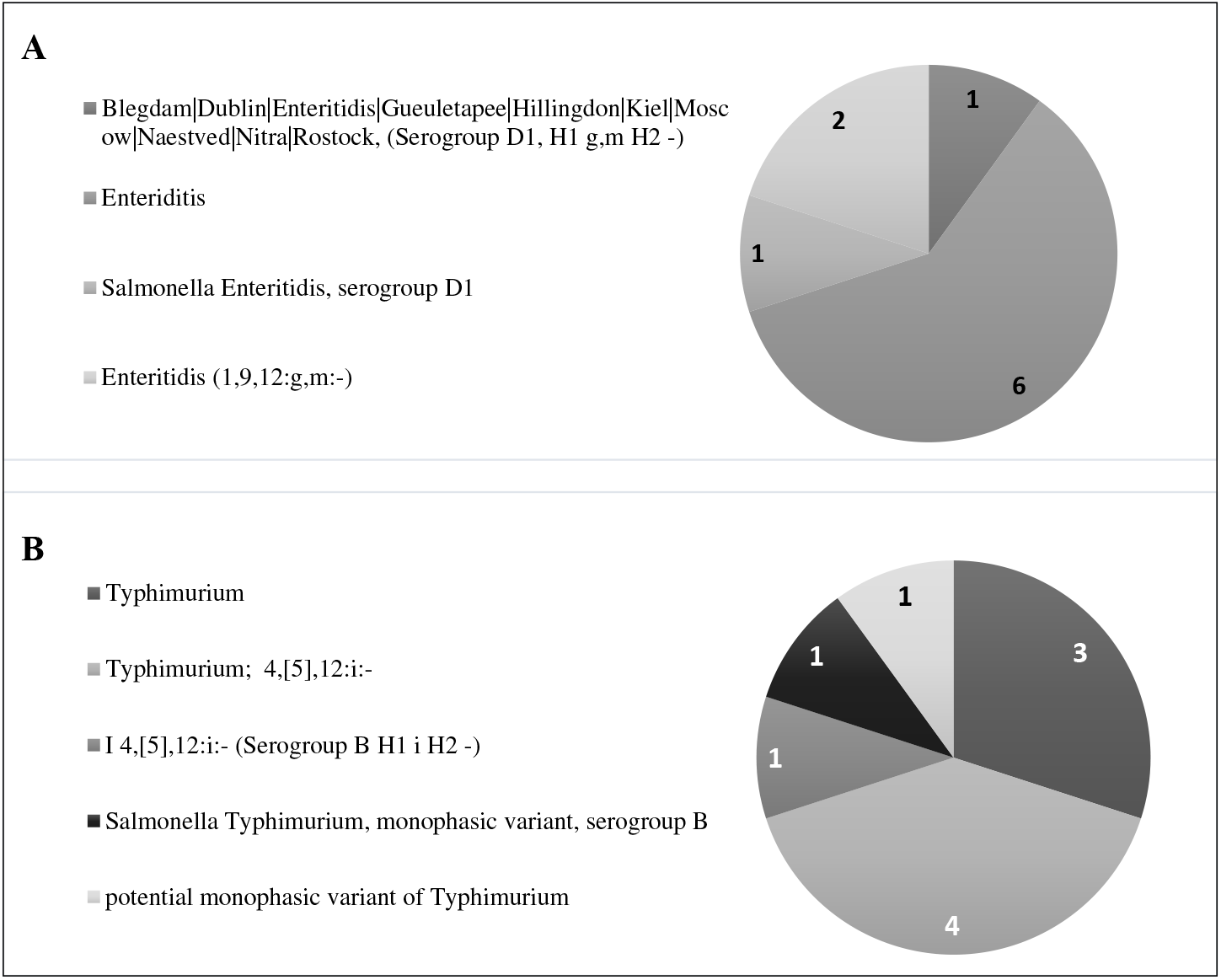
*In silico* serotype for (A) BSQAP001 and (B) BSQAP005, as reported by all participating laboratories (N=10) in the pilot PT.

**FIG 2:**
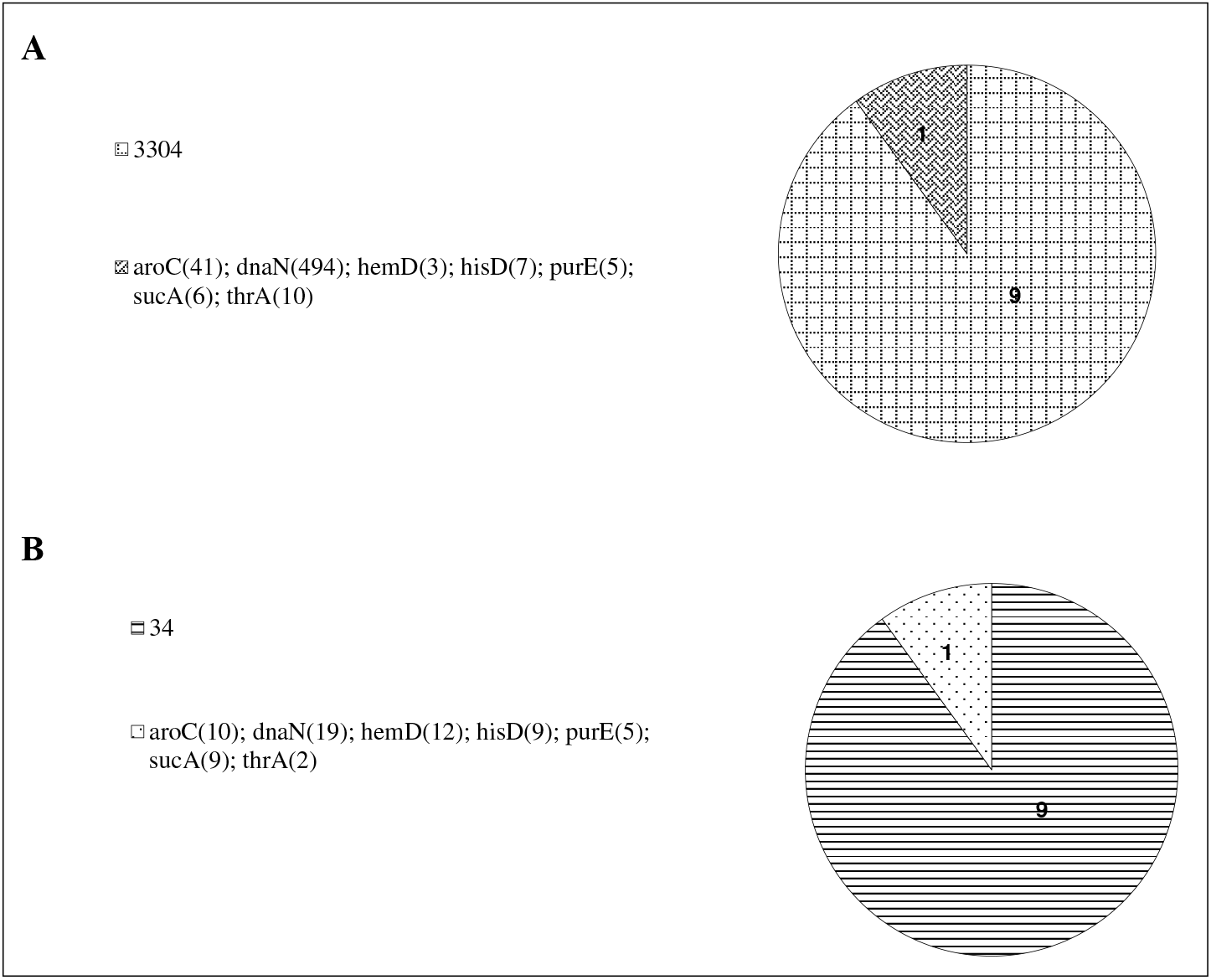
*In silico* MLST for (A) BSQAP001 and (B) BSQAP005, as reported by all participating laboratories (N=10) in the pilot PT.

**TABLE 3.** Antimicrobial resistance (AMR) genes harboured by different bacteria isolates were based on the sequencing data, as reported by the participating laboratories

| RCPAQAP SEQID | AMR genes | No. of participating laboratories that reported the AMR genes |
| --- | --- | --- |
| BSQAP001<sup>a</sup> | aac(6')-Iaa | 6/10 |
|  | mdf(A) | 4/10 |
|  | unspecified | 3/10 |
|  | no resistance genes detected | 1/10 |
| BSQAP005<sup>b</sup> | **tet(B)** | 10/10 |
|  | **sul2** | 10/10 |
|  | **blaTEM-1** | 9/10 |
|  | **aph(6)-Id** | 9/10 |
|  | **aph(3'')-Ib** | 8/10 |
|  | aac(6')-laa | 5/10 |
|  | mdf(A) | 3/10 |
|  | strB_1 | 1/10 |
|  | strA_4 | 1/10 |

<sup>a</sup> The bioinformatics analysis performed by the Bioinformatics Working Group, CDGN on all raw FASTQ data set submitted by participating laboratories for BSQAP001 found no complete or partial matches of AMR genes using X identity/Y coverage
<sup>b</sup> The bioinformatics analysis performed by the Bioinformatics Working Group, CDGN on all raw FASTQ data set submitted by participating laboratories for BSQAP005 identified five complete or partial matches of AMR genes (**in bold**)

### Protocol used in generating the WGS data

All laboratories used a commercial kit to prepare the sequencing library; six laboratories used the Illumina Nextera XT DNA Library Preparation Kit, while the remaining laboratories used the Illumina DNA Flex Library Prep Kit. All laboratories performed paired-end sequencing on the Illumina platform, employing the NextSeq (n=5), MiSeq (n=4) and MiniSeq (n=1); read lengths employed on the MiSeq were 150bp (n=2) and 300bp (n=2) and 75bp on the MiniSeq. Additional information on the protocol used to extract the DNA for sequencing is listed in Table S7, while the summary of the bioinformatics analysis software and tools used by laboratories is listed in Table S8.

### Assessment of laboratories performance

The raw FASTQ data submitted by the laboratories were analysed, and the individual and overall performance of laboratories in sequencing both samples was evaluated based on the observed values for the quality metrics (Table 4). All laboratories passed the assessment for all quality metrics, except the inferred genome coverage, whereby eight and six laboratories passed the assessment for BSQAP001 and BSQAP005, respectively. Two laboratories sequenced more than 1,000,000 bp nucleotides for NTC, and therefore failed the assessment for the total sequenced nucleotides (Fig. 3).

**FIG 3:**
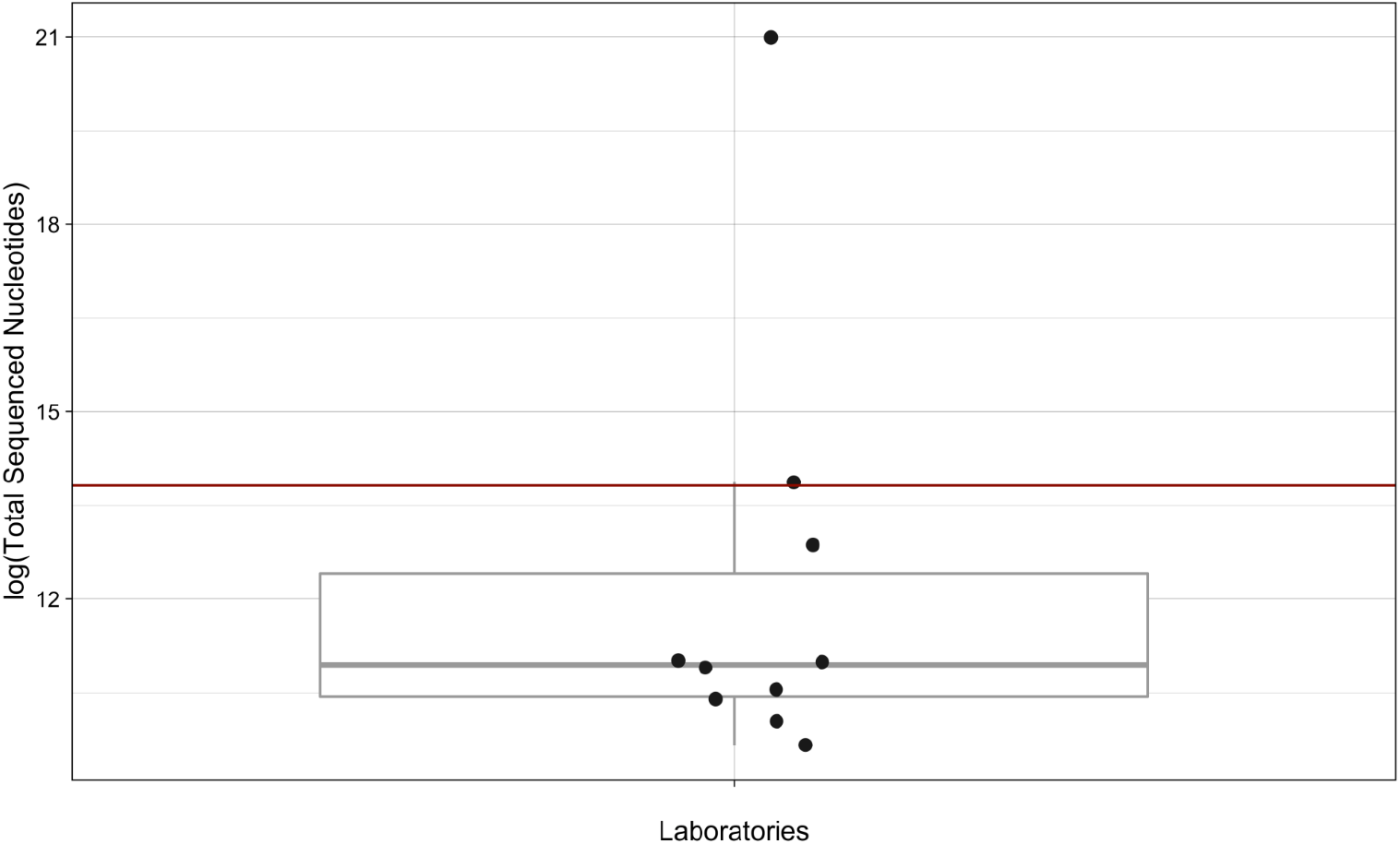
Comparative data of the NTC based on the total sequenced nucleotides performed by all the participating laboratories (N=10). The interquartile range is represented by the box, and the median is represented by the horizontal line, while the red line indicates 1,000,000 bp.

**TABLE 4.** The observed value of quality metrics based on the raw FASTQ data set submitted by each participating laboratory for; A) BSQAP001, B) BSQAP005, and the assessment of their overall performance

| Quality metrics | Participating laboratory<sup>a</sup> |  |  |  |  |  |  |  |  |  | Overall performance<sup>b</sup> |
| --- | --- | --- | --- | --- | --- | --- | --- | --- | --- | --- | --- |
|  | 1 | 2 | 3 | 4 | 5 | 6 | 7 | 8 | 9 | 10 |  |
| Average Q-Score | P | P | P | P | P | P | P | P | P | P | 10/10 |
| Inferred genome coverage | P | P | P | P | P | F | P | F | P | P | 8/10 |
| Inferred organism | P | P | P | P | P | P | P | P | P | P | 10/10 |
| Assembly genome size | P | P | P | P | P | P | P | P | P | P | 10/10 |
| Inferred MLST scheme | P | P | P | P | P | P | P | P | P | P | 10/10 |
| Inferred MLST ST | P | P | P | P | P | P | P | P | P | P | 10/10 |

B. BSQAP005
| Quality metrics | Participating laboratory<sup>a</sup> |  |  |  |  |  |  |  |  |  | Overall performance<sup>b</sup> |
| --- | --- | --- | --- | --- | --- | --- | --- | --- | --- | --- | --- |
|  | 1 | 2 | 3 | 4 | 5 | 6 | 7 | 8 | 9 | 10 |  |
| Average Q-Score | P | P | P | P | P | P | P | P | P | P | 10/10 |
| Inferred genome coverage | P | F | P | P | F | F | P | F | P | P | 6/10 |
| Inferred organism | P | P | P | P | P | P | P | P | P | P | 10/10 |
| Assembly genome size | P | P | P | P | P | P | P | P | P | P | 10/10 |
| Inferred MLST scheme | P | P | P | P | P | P | P | P | P | P | 10/10 |
| Inferred MLST ST | P | P | P | P | P | P | P | P | P | P | 10/10 |

### Comparison of the FASTQ data set submitted by laboratories

By analysing all participant data with a single pipeline, we control for the bioinformatics and are able to get a clear assessment of the effect of variability in sequencing approaches on downstream analyses. The comparative figure for both samples was summarised in Fig. 4, and the raw data is available in Fig S1. Overall, the results show that most laboratories sequenced to about the same effort, as suggested by the distribution of Z-scores for total reads and total sequenced nucleotides (most data are within one standard deviation from the mean). The exception is one laboratory that produced far more data than any other laboratory for both samples. While the average Q score was roughly bimodal, the percent GC also showed a bimodal distribution for each of the samples.

**FIG 4:**
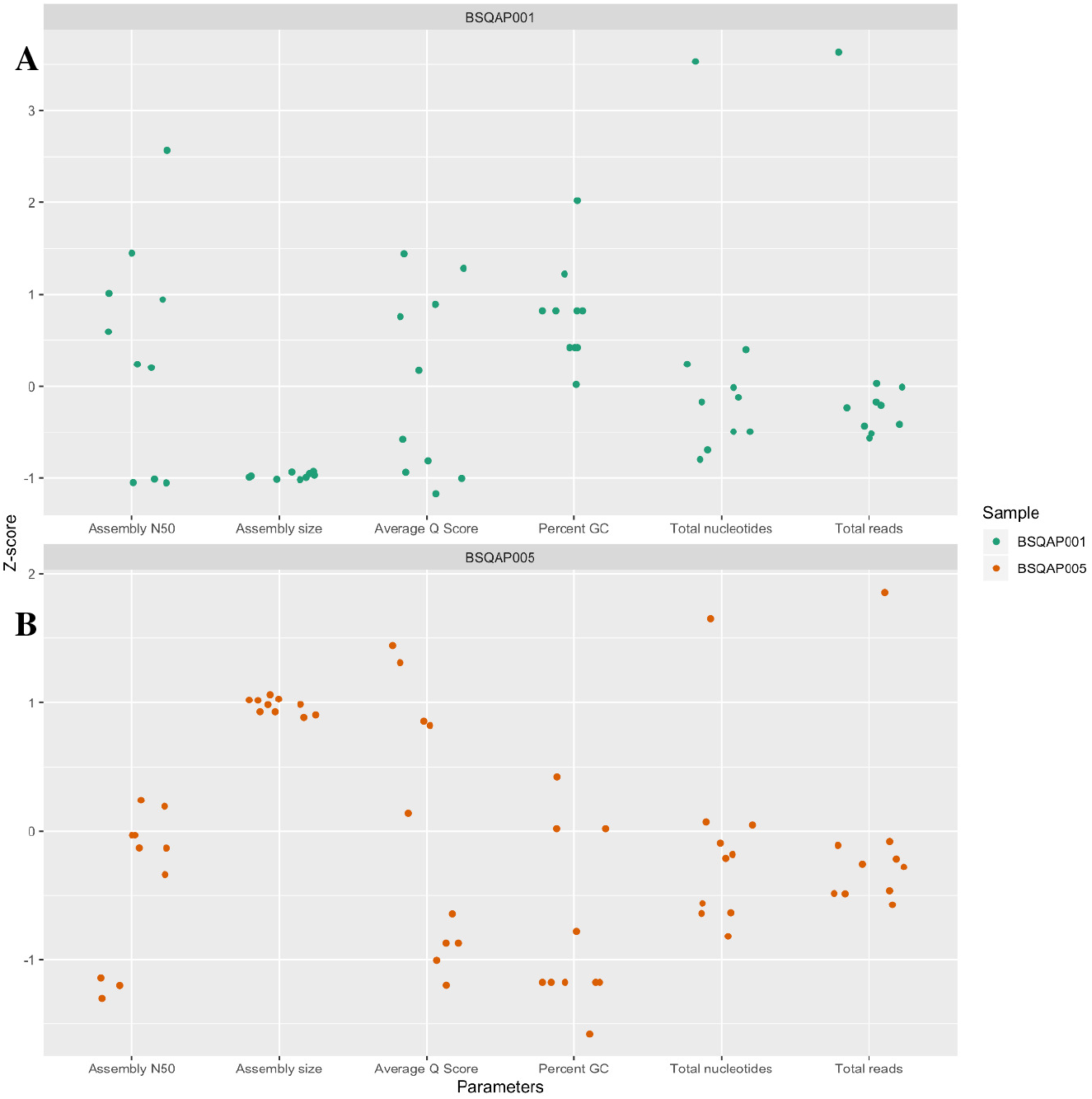
Comparison of the assessment of the effect of variability in sequencing approaches on downstream analyses data for (A) BSQAP001 and (B) BSQAP005, based on the analysis performed on all participant data (N=10) using a single pipeline.

The total assembly size for each sample was consistent within samples, and not unexpectedly distinct between samples. Despite consistent assembly sizes within samples, there was considerable variation, with Z-scores spanning almost 3.5 standard deviations in the case of sample BSQAP001.

## DISCUSSION

The effective analysis of WGS data for infectious diseases requires a multidisciplinary team with technical and bioinformatics skills, as well as biological, epidemiological and clinical knowledge. The intricate multistep processing of WGS outputs generated by different sequencing platforms begins with sequence assembly or reference-based mapping and ends with simultaneous comparisons of multiple genomes and data visualisation [23]. Apart from processes such as library preparation and quality assessment of sequences produced, it also requires sequence reads validation. The validation involves processing of sequence reads with software tools or assembled into workflows and pipelines. Subsequently, the results obtained from the bioinformatics analysis for pathogen identification are evaluated post-analytically, including making correlations between bioinformatics results with clinical and epidemiological patient information.

The aim of this study was to establish a workflow that allows assessment of a laboratory’s proficiency to perform WGS on bacterial isolates belonging to the genus *Salmonella*. This included defining minimum quality matrices, building a bioinformatics pipeline to analyse laboratories results, and developing a PT program with appropriately tested specimens and results reporting facilities. In this pilot PT, the capacity of laboratories to perform WGS was assessed initially on the correct organism identification, the determination of MLST, and the detection of AMR genes. The data submitted by all laboratories were analysed with a single pipeline, to ensure that the bioinformatics were controlled and the effect of variability in sequencing approaches on downstream analyses was clearly assessed. Essentially, participating laboratories were capable in generating high-quality data, either meeting or exceeding the minimum requirements. In some cases, there were laboratories that far exceeded the minimum requirements and could potentially save money by reducing their sequencing effort. While all laboratories determined the correct organism, serotype and sequence type, we observed some variations in the format of data entry and terminology. There were significant variations observed in the reporting of presence or absence of AMR genes by all the laboratories. This was likely due to the high variation in assembly, which may impact on the post sequence analysis. Also, the wide-ranging approaches used in the genome-wide analyses, the gene detection software and database used, as well as the thresholds applied to determine the presence or absence of AMR genes may have affected the report of the AMR genes for both samples. The results from this study corroborate those of observed in a similar study [24].

The detection of genomic variants will be affected by specimen preparation, sequencing methods, run throughput, and the amount of sequencing data sufficient for pathogen characterisation (genome coverage), with the choice of technology and coverage being particularly important. Prior to the sequencing step, DNA libraries need to be constructed, usually with protocols that are significantly variable. However, this critical step can have significant influence in downstream analyses, thus selecting the most cost-effective library construction approach is crucial to ensuring the highest quality data is available for inferences. To date, the data quality produced by different library methods has only been addressed in few studies. While the data quality is highly dependent on the purity of DNA sample submitted for sequencing and its GC-content [25], the choice of library preparation method is also known to influence the level of sequencing quality. Given that libraries were also run on different instruments which may introduce own biases, the process of comparison therefore will become too complex. While this study did not compare the quality of data based on the choice of library preparation method used, the performance of laboratories in generating the WGS data set in the future PT can potentially be measured using this aspect of data generation, if the right approaches are applied. In this pilot PT, commercial kits were used by all participating laboratories in the library preparation for sequencing. Kits that are based on the enzymatic shearing of DNA, like the Illumina Nextera XT used by most of the laboratories in this pilot PT may result in less uniform data quality and missing more specific genomic sequence motifs (coverage bias), compared to the mechanical method in the Illumina TruSeq kit, as reported in a study performed on the GC-rich *Mycobacterium tuberculosis* genome [25]. The remainder of the laboratories here used the Illumina DNA Flex kit, a robust transposome bead-based method which has been reported to give a consistent yield and fragment size, and less sequence bias [26]. It remains a debate whether the process of generating the library using either the Illumina Nextera XT or the Illumina DNA Flex kit is significantly different, given that both kits are transposome-based. The only difference is in the method that is applied to normalise the sample and in the delivery of the transposome (in solution with Nextera XT versus on a bead with Flex), and this could also be an aspect of WGS data generation to be assessed in the future PT. A recent study evaluating the commercial library preparation kits have showed that the data quality from libraries made with Illumina DNA Flex kit were superior to those from Nextera XT. This was due to decreased sensitivity to variable GC-content, therefore an almost uniform distribution of read-depth and minimal low coverage regions, resulting in a more complete representation of the genome [27]. However, it has also been reported that more adapter dimer products are often generated, introducing contamination when sample input is limiting, as a results from suboptimal performance of the bead-based method [28], and potentially due to bead-manufacturing process (personal communication).

Several technologies have been used over the past decade for WGS: Ion Torrent PGM, PacBio, Roche 454, and more recently Oxford Nanopore Technologies. However, for bacterial genomics, Illumina machines are the most often used platform for generating WGS data (>90% at the Short Read Archive) [27]. Mirroring international practises, all participating laboratories of this PT used Illumina sequencing platforms (either the NextSeq, MiSeq or MiniSeq platforms).

The international public health community is still developing standards for pathogen identification, typing, microbial resistance detection and surveillance using WGS data (see PHA4GE initiative at https://pha4ge.github.io/). There is a need for these standards to be implemented to help harmonize the analysis and the consistency in the interpretation of WGS sequence data [29], [30]. It is also critical due to the various potential sources for error in sequencing and variant-calling processes [31], [32]. To optimise the quality of the data used to generate variant calls, many of the recommendations and best practises used are shared between human and microbial genomic NGS assays [33], [34]. This includes the recommendation to minimise amplification steps in library preparation (if applicable), use of paired-end sequencing, removal of duplicate reads, realignment around insertions and deletions, and recalibration of base Q-scores. In the absence of definitive quality metrics to measure the capacity of the participating laboratories in performing the WGS, we have employed six different quality metrics in this pilot PT, following the discussion within the CDGN Bioinformatics Working Group. Among the quality metrics that were assessed, only a minority of the participating laboratories failed to sequence to the required genome coverage depth of at least 40X for both samples. Among them, one laboratory particularly had a very low coverage (less than 20X) for both samples, potentially resulting in decreased sensitivity in their WGS data. Sufficient sequence coverage is vital for the accurate identification of genetic variants, the association between higher coverage levels and the steady increase in the sensitivity in WGS data, as previously reported [35]. The sequencing data from the NTC are useful not only to identify cross-contamination but also to detect any potential contaminants in buffers and reagents. The role of NTC in WGS is still being explored and best-practices are still being in development. While most published studies on the use of WGS for diagnostics and WGS PT did not incorporate a NTC analysis, we are proponents that an NTC is absolutely required for high-quality data in a public health and clinical setting, as it is also part of the requirement for accreditation. In most cases, however, a failed NTC will not cause a whole run to be failed. That approach would be too costly. Instead, we propose that a failed NTC triggers a more in-depth analysis of each sample in the run. Samples found to have significant evidence of contamination should be failed. Although, the levels of contamination deemed to be significant remain an open question. In this pilot PT, only two laboratories failed the assessment for the total sequenced nucleotides and will be required to further investigate and use this information to decide on what samples should or should not be failed. While the Z-score comparative data between the participating laboratories did not need to cluster together, ideally the results would cluster together for consistency across the analysis performed by the laboratories. In this pilot PT, the deviations in assembly N50 may suggest that the quality of the genome assembly performed by different laboratories was varying. This also further suggests fragmentation variation to some degree, for instance, due to contamination which may potentially affect the downstream gene detection. Some variation in assembly was also observed from the comparative data, which may impact on post sequencing analysis. Different versions of bioinformatics tools (e.g. *abricate*), choices of reference database or a different threshold applied in deducing the presence/absence of genes, may in turn impact on results. Other downstream variations in analysis, for instance, the average Q-score and percent GC were within the acceptable range. Q-score is an indication of the probability of error, whereby higher Q-scores indicate a smaller probability of error and lower Q-scores can result in a significant portion of the reads being unusable, which in turn may also lead to increased false positive variant calls, resulting in inaccurate conclusions. The average Q score was roughly bimodal, and we believe this largely reflects the instrument used (MiSeq vs NextSeq). Significant deviations in the GC-content and assembly size may be indicative of contamination from an unrelated species or source with a different GC profile. While the data on total nucleotides and total reads clustered closer together for most participating laboratories, these were also a reflection on whether the laboratories have performed sufficient sequencing to yield the required sequence information for analysis (not over or under sequencing). Total nucleotides sequenced is a better comparison measure of the quality of the sequence, compared to the total reads, as the read length can vary depending on the library preparation. The consistency in the total nucleotides will affect downstream applications; while excessive coverage could increase the false rate in the single nucleotide polymorphism (SNP) detection and assembly and the cost to sequence per isolate; inadequate coverage may result in missing out on SNPs and issues with assemblies, and subsequently, gene detection.

In this pilot PT, we observed some minimal variation in methods employed to generate and analyse the data, as well as in terms of quality control criteria, as expected in this early stage of the transition to genomics. Almost all participating laboratories used *kraken, mlst*, and *abricate* for sample ID, MLST and AMR, respectively, and most if not all have applied similar workflows, (i.e. Nextera XT in preparing the library, Illumina sequencer in sequencing the samples and Qiagen-based DNA extraction system) resulting in no extensive variation in bioinformatics tools and workflows for infectious agent analysis of WGS data. The only variation is likely in the versions used and the setting applied, therefore highlighting that harmonization is not just dependent on choice of tool used, but version control. With an appropriate validation approach for the choice of tool and version, this variation may not affect the quality of the WGS data or in measuring the capability of clinical laboratories in performing WGS (both wet and dry-laboratory components). There will never be a standard procedure for this analysis, which therefore makes the comparison of such analyses complex. With an abundance of tools continually being developed, refined and packaged together as bioinformatics pipelines, it is expected that the variation in the bioinformatics approaches used in the microbiology community will remain highly variable [36]. Another challenge is in defining the ‘truth’ of the sample set used in a PT. Where there is high confidence in the truth of the WGS PT results and clear guidance on the settings to be used to define the truth, laboratories will then able to use that to calibrate their testing and results via a PT program.

In summary, the introduction of a WGS PT program such as this pilot PT will facilitate harmonisation of genomics implementation, including standardisation of bioinformatics workflow, thereby making comparisons across jurisdictions more robust, reduce ambiguity and bolster confidence in the data generated and the process employed. Lessons learned from this pilot PT will help inform the development of future WGS PT programs for infectious diseases and create a consistent mechanism in conducting continuous PT to guarantee the quality of the WGS data generated from clinical and public health laboratories. The pilot PT offered in this study has been well received, due to the limitation in the availability of an external PT program for WGS of infectious agents for Australian laboratories. It demonstrated the capacity and protocol used by clinical and public health laboratories around Australia in performing WGS and analysing WGS data. In the future, laboratories are expected to create a balance between the characterisation of pathogen genome, the throughput of the instrument, and the accuracy of variant-calling algorithms.

## ACKNOWLEDGEMENT

RCPAQAP Biosecurity is funded by the Australian Government Department of Health.

We thank the Bioinformatics and Genomics Implementation Working Groups within the Communicable Diseases Genomics Network (CDGN), an Expert Reference Panel under the Public Health Laboratory Network (PHLN), Australia, in assisting the development of the 2018 Pilot PT program for WGS of infectious agents.

## REFERENCES

1. Long, S.W., et al., A genomic day in the life of a clinical microbiology laboratory. J Clin Microbiol, 2013. 51(4): p. 1272–7.

2. Didelot, X., et al., Transforming clinical microbiology with bacterial genome sequencing. Nat Rev Genet, 2012. 13(9): p. 601–612.

3. Kalman, L.V., et al., Current landscape and new paradigms of proficiency testing and external quality assessment for molecular genetics. Arch Pathol Lab Med, 2013. 137(7): p. 983–8.

4. Moran-Gilad, J., et al., Proficiency testing for bacterial whole genome sequencing: an end-user survey of current capabilities, requirements and priorities. BMC Infect Dis, 2015. 15: p. 174.

5. Timme, R.E., et al., GenomeTrakr proficiency testing for foodborne pathogen surveillance: an exercise from 2015. Microb Genom, 2018. 4(7).

6. Allard, M.W., et al., Practical Value of Food Pathogen Traceability through Building a Whole-Genome Sequencing Network and Database. J Clin Microbiol, 2016. 54(8): p. 1975–83.

7. Brinkmann, A., et al., Proficiency Testing of Virus Diagnostics Based on Bioinformatics Analysis of Simulated In Silico High-Throughput Sequencing Data Sets. J Clin Microbiol, 2019. 57(8).

8. Hutchins, R.J., et al., Practical Guidance to Implementing Quality Management Systems in Public Health Laboratories Performing Next-Generation Sequencing: Personnel, Equipment, and Process Management (Phase 1). Journal of Clinical Microbiology, 2019. 57.

9. Koster, J. and S. Rahmann, Snakemake-a scalable bioinformatics workflow engine. Bioinformatics, 2018. 34(20): p. 3600.

10. Kurtzer, G.M., V. Sochat, and M.W. Bauer, Singularity: Scientific containers for mobility of compute. PLoS One, 2017. 12(5): p. e0177459.

11. Li, H. Seqtk: Toolkit for Processing Sequences in FASTA/Q Formats. 2018; Available from: https://github.com/lh3/seqtk.

12. Bolger, A.M., M. Lohse, and B. Usadel, Trimmomatic: a flexible trimmer for Illumina sequence data. Bioinformatics, 2014. 30(15): p. 2114–20.

13. Wood, D.E. and S.L. Salzberg, Kraken: ultrafast metagenomic sequence classification using exact alignments. Genome Biol, 2014. 15(3): p. R46.

14. Ondov, B.D., et al., Mash: fast genome and metagenome distance estimation using MinHash. Genome Biol, 2016. 17(1): p. 132.

15. Seemann, T., et al. Shovill: Faster SPAdes Assembly of Illumina Reads. 2018; Available from: https://github.com/tseemann/shovill.

16. Bankevich, A., et al., SPAdes: a new genome assembly algorithm and its applications to single-cell sequencing. J Comput Biol, 2012. 19(5): p. 455–77.

17. Gurevich, A., et al., QUAST: quality assessment tool for genome assemblies. Bioinformatics, 2013. 29(8): p. 1072–5.

18. Seemann, T. Mlst: Scan Contig Files Against PubMLST Typing Schemes. 2018; Available from: https://github.com/tseemann/mlst/.

19. Jolley, K.A., J.E. Bray, and M.C.J. Maiden, Open-access bacterial population genomics: BIGSdb software, the PubMLST.org website and their applications. Wellcome Open Res, 2018. 3: p. 124.

20. Seemann, T. ABRicate: Mass Screening of Contigs for Antimicrobial and Virulence Genes. 2018; Available from: https://github.com/tseemann/abricate.

21. NCBI, NCBI AMR Reference Gene Database.

22. Yoshida, C.E., et al., The Salmonella In Silico Typing Resource (SISTR): An Open Web-Accessible Tool for Rapidly Typing and Subtyping Draft Salmonella Genome Assemblies. PLoS One, 2016. 11(1): p. e0147101.

23. Carrico, J.A., et al., A primer on microbial bioinformatics for nonbioinformaticians. Clin Microbiol Infect, 2018. 24(4): p. 342–349.

24. Doyle, R.M., et al., Discordant bioinformatic predictions of antimicrobial resistance from whole-genome sequencing data of bacterial isolates: An inter-laboratory study. bioRxiv, 2019.

25. Tyler, A.D., et al., Comparison of Sample Preparation Methods Used for the Next-Generation Sequencing of Mycobacterium tuberculosis. PLoS One, 2016. 11(2): p. e0148676.

26. Bruinsma, S., et al., Bead-linked transposomes enable a normalization-free workflow for NGS library preparation. BMC Genomics, 2018. 19(1): p. 722.

27. Seth-Smith, H.M.B., et al., Evaluation of Rapid Library Preparation Protocols for Whole Genome Sequencing Based Outbreak Investigation. Frontiers in Public Health, 2019. 7(241).

28. Head, S.R., et al., Library construction for next-generation sequencing: overviews and challenges. BioTechniques, 2014. 56(2): p. 61–passim.

29. Gwinn, M., D. MacCannell, and G.L. Armstrong, Next-Generation Sequencing of Infectious Pathogens. JAMA, 2019. 321(9): p. 893–894.

30. Williamson, D.A., et al., The importance of public health genomics for ensuring health security for Australia. Med J Aust, 2019. 210(7): p. 295–297.e1.

31. Bush, S.J., et al., Genomic diversity affects the accuracy of bacterial SNP calling pipelines. bioRxiv, 2019.

32. Yoshimura, D., et al., Evaluation of SNP calling methods for closely related bacterial isolates and a novel high-accuracy pipeline: BactSNP. Microbial Genomics, 2019. 5(5).

33. Olson, N.D., et al., Best practices for evaluating single nucleotide variant calling methods for microbial genomics. Front Genet, 2015. 6: p. 235.

34. Gargis, A.S., et al., Good laboratory practice for clinical next-generation sequencing informatics pipelines. Nat Biotechnol, 2015. 33(7): p. 689–93.

35. Chiara, M. and G. Pavesi, Evaluation of Quality Assessment Protocols for High Throughput Genome Resequencing Data. Front Genet, 2017. 8: p. 94.

36. Wyres, K.L., et al., WGS Analysis and Interpretation in Clinical and Public Health Microbiology Laboratories: What Are the Requirements and How Do Existing Tools Compare? Pathogens, 2014. 3(2): p. 437–58.

